# Data-driven motif discovery in biological neural networks

**DOI:** 10.1101/2023.10.16.562590

**Authors:** Jordan K. Matelsky, Michael S. Robinette, Brock Wester, William R. Gray-Roncal, Erik C. Johnson, Elizabeth P. Reilly

## Abstract

Data from a variety of domains are represented as graphs, including social networks, transportation networks, computer networks, and biological networks. A key question spans these domains: are there meaningful repeated subgraphs, or motifs, within the structure of these larger networks? This is a particularly relevant problem when searching for repeated neural circuits in networks of biological neurons, as the field now regularly produces large brain connectivity maps of neurons and synapses, or connectomes. Given acquisition costs, however, these neuron-synapse connectivity maps are mostly one-of-a-kind. With current graph analysis techniques, it is very challenging to discover new “interesting” subgraphs *a priori* given small sample sizes of host graphs. Another challenge is that for even relatively modest graph sizes, an exhaustive search of all possible subgraphs is computationally intractable. For these reasons, motif discovery in biological graphs remains an unsolved challenge in the field. In this work, we present a motif discovery approach that can derive a list of undirected or directed motifs, with occurrence counts which are statistically significant compared to randomized graphs, from a single graph example. We first address common pitfalls in the current most common approaches when testing for motif statistical significance, and outline a strategy to ameliorate this problem with improved graph randomization techniques. We then propose a progressive-refinement approach for motif discovery, which addresses issues of computational cost. We demonstrate that our sampling correction technique allows for significance testing of target motifs while highlighting misleading conclusions from standard random graph models. Finally, we share our reference implementation, which is available as an open-source Python package, and demonstrate real-world preliminary results on the *C. elegans* connectome and the ellipsoid body of the *Drosophila melanogaster* fruit fly connectome.

## Introduction

Many research domains use graph data structures to organize complex, interconnected data. Social networks are commonly represented as users (vertices) connected by “friendships” or other relationships (edges). Biological neuronal networks are now commonly represented as *nanoscale connectomes*, where vertices represent neurons and edges represent directional synapses between them [1, 2, 3]. In all such domains, one common research question is to identify important repeating subgraphs, or *motifs*. In a social media network, this might be in pursuit of identifying friend-groups, detecting artificially inflated botnets, or isolating terrorist groups [4]. In the field of connectomics, neuroscience research has long posited that desirable properties of the brain as a learning machine (such as energy efficiency or active learning) may be due to the brain’s deployment of repeating computational substructures [5]. As emerging comparative connectomics datasets across disease states or development [6] emerge, studying changes amongst these repeated structures could highlight critical underlying structural changes. In addition, artificial neural network (ANN) research has already greatly benefited from neuro-inspired structural circuits such as convolution and recurrence [7]. Ongoing neuroscience-inspired machine learning efforts [8, 9] may benefit not only from an understanding of the function of neural circuits, but also key repeated structural components in the brain at different spatial scales[10, 11, 12, 13].

Historically, finding these motifs in the brain has been a labor-intensive process: neuroscientists usually posit the structure of a motif based on preliminary structural or functional data, and then have to verify or refute the hypothesis by examining the brain’s connectome through manual inspection [14]. More recent techniques [10, 11, 12] have enabled optimized, high-speed search of the connectome for a given motif. But despite these advances, it is still very challenging to discover new motifs *a priori*. This is due in part to the fact that neuronal network connectomes can be very expensive and time-consuming to generate, requiring expert personnel and specialized equipment. Furthermore, connectomes are often one-of-a-kind, meaning that traditional statistical models are challenging or impossible to implement. Finally, exhaustive subgraph matching or counting can be computationally intractable for even relatively small connectomes [15, 16].

Thus, the challenge for a motif discovery pipeline is bipartite: First, such a tool must operate in a *N* = 1 regime. In other words, it must be able to operate on a single host network, rather than a population of real-world samples. And second, it must be able to rapidly downsize the field of possible motifs so that the number of exhaustive subgraph matches performed is as small as possible.

Here, we outline our greedy-algorithm approach for data-driven motif discovery, validate it on artificial as well as biological networks, and show that our approach can discover motifs through an automated, data-driven process with no prior knowledge of the connectome. Finally, we comment on the motifs discovered through this pipeline, and validate our results.

## Background

Several critical advancements have been made in the analysis of biological neural networks, including scalable subgraph isomorphism search [10], identification of hierarchical structure [17], and network motif enumeration [18] including edge coloring [19]. In addition, there are several existing motif discovery methods, many of which have been applied to problems in bioinformatics. Here, we consider methods that seek to identify subgraphs that occur more frequently than expected in a relevant random graph model. Note that these methods might not find every functionally significant subgraph. For instance, there may be functionally significant subgraphs that don’t occur very frequently. However, given their placement within the overall network or other network characteristics, they may still have an interesting computational role. Therefore, a full discovery process on a network of interest should utilize multiple approaches for identifying significant structure and function within the network, including motif discovery approaches.

As seen in [20], structural motif discovery methods are characterized by several properties. Some methods are motif centric, meaning they start with a list of candidate motifs, and others are network centric, meaning they simply count all subgraphs of a specific size within the network. This has the advantage of only searching for subgraphs that are known to exist in the network. However, through our experience thus far with noisy real world networks, nearly all small subgraphs *do* tend to exist in even moderately-dense networks, which result in little difference between these two approaches. Methods also differ in the way they determine significance. Some approaches include threshold testing and significance testing. Different metrics may be used for these tests. Some approaches use sampling methods as an alternative to ‘exact enumeration’ to improve computational speed. There are also various subgraph isomorphism algorithms that can be leveraged. And, of course, there are a number of additional heuristics that can be used to improve computational feasibility. Some of the popular approaches include MAVisto, MFinder, FANMOD, and MODA [21, 22, 23, 24, 25], as well as our more recent contributions, DotMotif and GrandIso [10] and their cloud-scale equivalents [15].

The challenge of *N* = 1 network analysis has been addressed in the past through various techniques. Most common among these techniques are the generation of a population of similar networks, using random graph techniques. The simplest — and most common — random graph models for biological network analysis [26, 10] are the Erdős-Rényi model, in which every edge is independent and equally likely [27]; the Geometric random graph model, in which vertices are independently and uniformly embedded in the d-dimensional Euclidean space R^*d*^ and edges are formed between pairs of vertices whose Euclidean distance is below a given threshold ∥*v*_*i*_ *− v*_*j*_ ∥ *≤r*, where ∥ *·* ∥ represents the Euclidean distance and *r* is the given threshold [28]; the Barabási-Albert graph, which is generated by starting with a growing network *G* = (*V, E*), with each new node *v*_*i*_ added at time step *t* connecting to existing nodes *v*_*j*_ with probability proportional to their degree *k*_*j*_, resulting in a scale-free network where the degree distribution follows a power-law *P* (*k*) *∝ k*^*−γ*^ [29]; and the Watts-Strogatz graph which starts with a regular ring lattice of *n* nodes with each node connected to its *k* nearest neighbors, and then rewires each edge independently with probability *p*, resulting in a graph with high clustering and small average path length [30].

Each of these models have certain strengths (low parametrization dimension; simplicity of model) but all fail to represent the local topology of the host graph to which they are calibrated in a way that roughly conserves subgraph count [10]. This means that using these random graph models as a null model against which to compare motif counts of a connectome will yield misleading results. In contrast, the *X*-swap degree-preserving operation (**Fig. 1B-C**) by which pairs of edges from a starting graph are swapped repeatedly, conserves certain desirable properties of a network (such as degree distribution of each vertex) [31]. However, the traditional *X*-swap operation has certain side-effects that mean edge-swapping is neither ergodic nor unbiased [32]. Controlling for these side-effects as described below, it is this algorithm on which we base our motif discovery pipeline.

**Figure 1.**
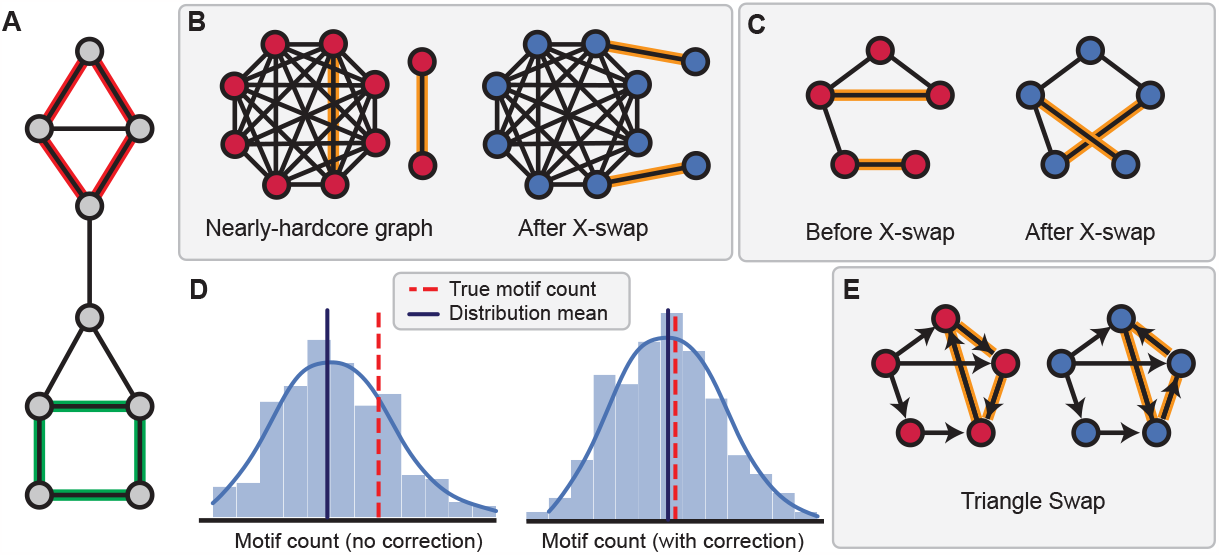
Illustration of X-swap and triangle swap operations. **A**. There are two valid subgraph monomorphisms of the four-vertex cycle, one highlighted in red and one highlighted in green. The orange monomorphism is *not* a valid isomorphism due to the horizontal edge. **B-C**. The X-swap operation can change the number of motifs without affecting degree distribution. In this example, the highlighted edges in the unmodified graph (**a**) are swapped. Although the global graph degree sequence, as well as the degree of each node, remains the same, the modified graph in **C** has a new 4-cycle subgraph (highlighted in red) where it did not before exist. Likewise, the X-swap operation can *remove* this cycle from the modified graph if run a second time. **D** Demonstration of the corrected swap operation on the motif count distribution, which is necessary for the distribution to be unbiased. **E** The triangle swap operation before (left) and after (right), which is used in the corrected algorithm.

## Definitions

A **graph** is a pair *G* = (*V, E*), where *V* is a set of vertices and 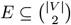 is a set of edges connecting those vertices. A **subgraph isomorphism** is a mapping of the edges of some graph *G* = (*V*_*G*_, *E*_*G*_) (the “query” graph) to a subset of the edges of another graph *H* = (*V*_*H*_, *E*_*H*_) (the “host” graph) subject to a bijective function *f* : *V*_*G*_ *→V*_*H*_ such that *u, v ⊆E*_*G*_ and *f* (*u*), *f* (*v*) ⊆ *E*_*H*_ . A **monomorphism** loosens this requirement from bijection to *injection*, meaning that the induced subgraph in *H* may include additional edges (**Figure 1A**). We define a *host* graph as the network inside of which we are searching for subgraph structure. We define a *query graph* or *query motif* as a subgraph structure that may or may not appear in the host graph. In this paper, we define motifs to be subgraphs of particular interest, perhaps due to expected existence based upon prior literature, or other distinguishing characteristics. Specifically, we will use the term *motif* to represent a discovered or putative subgraph that is over-or under-expressed compared to the expected count based upon our *X*-swap null model (see **A bias-corrected *X*-swap operation**).

## Results

### Motifs in *C. elegans*

The *C. elegans* nematode is the first organism for which a complete neural synaptic connectome was available [33]. Due to its relatively small size (only around three hundred neurons) and high stereotypy across individuals, *C. elegans* is a useful model organism for motif search [6]. The connectome network has a relatively low density 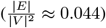 [10], which also makes this connectome particularly suitable for motif search on consumer compute hardware.

We ran our motif discovery pipeline on (directed) subgraph sizes up to | *V* | = 5. Our technique yields a cohort of significant motifs comprising both over-as well as under-expressed motifs (i.e., motifs that occur more frequently or less frequently, respectively, than we would expect to exist by chance). We render the top motifs discovered through our pipeline in **Fig. 2**.

**Figure 2.**
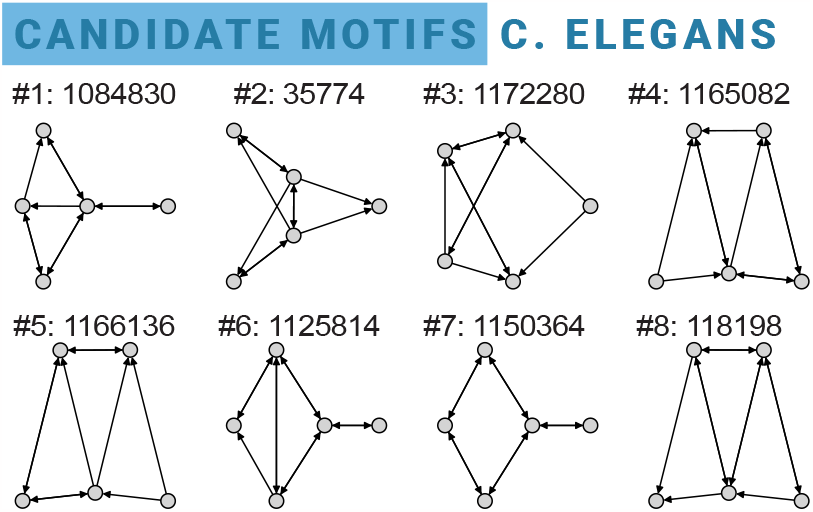
Top most-significant directed motifs discovered in the *C. elegans* synaptic connectome. These are the first eight motifs our pipeline recommends for further study in *C. elegans*. All of these networks are relatively high density and many include structures known to be relevant for computation, such as feedback loops and recurrence.

We can also begin to understand the cell-level composition connectome using motif analyses such as these. For example, **Fig. 3** shows how we can interrogate patterns at the cell level. Here motifs are characterized by the known cell types of *C. elegans* neurons. Further motif searches can interrogate specific cell types to refine the directed motif model.

**Figure 3.**
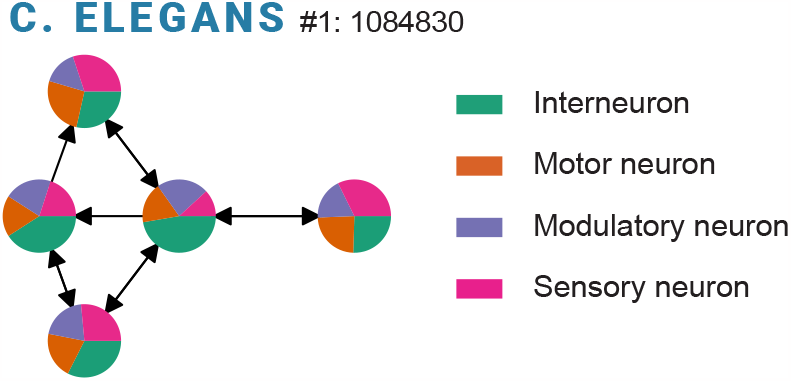
Cell-level analysis of an example motif in *C. elegans*. We render each vertex of the motif with a chart representing the neuron types in the host to which each vertex maps. From this display, we can gather that an interneuron “backbone” is visible (the horizontal axis is dominated by interneurons), and sensory neurons tend to be more sparsely connected than other types (sensory neurons tend to map to the more sparsely connected motif vertices, *i*.*e*., the rightmost vertex). More advanced analyses such as this one are possible with the analysis tools proposed here.

### Motifs in the *Drosophila* ellipsoid body

The ellipsoid body (EB) of the *Drosophila* fruit fly connectome is likewise regarded as fertile ground for investigations into neural circuits supporting memory, sensory integration, and navigation due to its well-understood function and well-characterized cell types. Here we study an excerpt of the *Hemibrain* [2, 3] connectome comprising the EPG, PEG, PENa, PENb, and Pintr cell types. This circuit has previously been studied and modeled computationally [34, 35] to investigate heading direction estimation in *Drosophila*.

We discover that the most significant motifs, according to our algorithm, tend to resemble a hub-and-spoke pattern, involving four vertices as a cycle with a fifth vertex as the shared hub (**Fig. 4**). It is noteworthy that the corresponding cycle and star graph do *not* appear in the set of most-significant motifs, suggesting that it is the cycle *in addition to* the hub-and-spoke that is repeated throughout the circuit.

**Figure 4.**
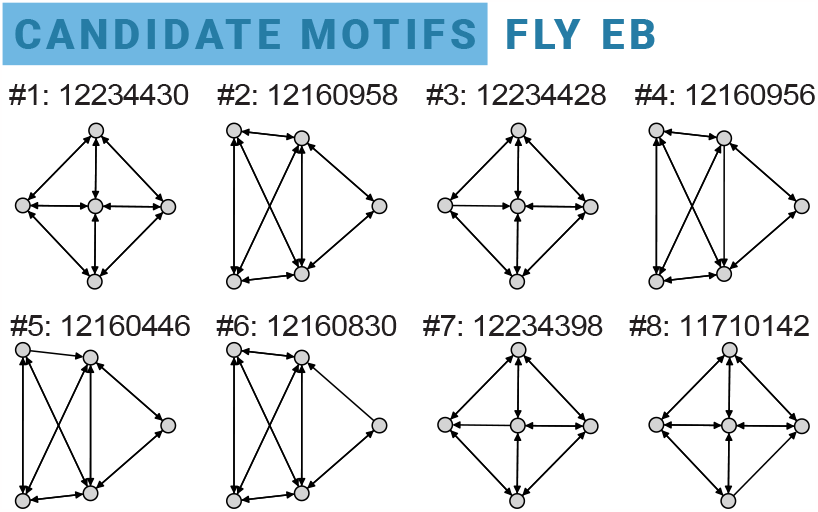
Top most-significant motifs discovered in the *Drosophila* ellipsoid body connectome. These eight motifs were determined to be the most significant in the ellipsoid body network. In contrast with the *C. elegans* motifs (**Fig. 2**), these motifs tend to almost always include a four-vertex cycle with a fifth central vertex connected to all of them, in a hub-and-spoke configuration.

## Discussion

Motif discovery on one-of-a-kind graphs is a sought-after capability in neuroscience and beyond. The technique proposed in this paper is an approach to addressing this problem, and there are opportunities to swap aspects of our pipeline for other software components: Subgraph isomorphism search, statistical testing, and the graph randomization or null-model generator are all modules that we anticipate being ripe for community contributions and further research.

Meanwhile, we believe there is much to be learned from the motifs discovered thus far in the *C*. elegans and *Drosophila* connectomes. That we discovered motifs with properties known to have biological relevance (e.g., recurrence, feedback and feedforward loops) is unsurprising. The next question is whether these motifs are conserved across individuals of the same species: Are these synaptic motif structures critical for the architecture of the animal, or are they a coincidence of more subtle and complex functional patterns? [36, 37] Or, as the truth is most likely some combination of both — *how* can we distinguish between “core” circuitry and periphery — and is the distinction even biologically important?

The question of the functional and dynamical implications of structural motif must be a major consideration for future work. This prospect, however, may bring with it some considerable advantages: Recent work has shown that attractor dynamics may correlate closely with certain structural motifs [38, 39], simultaneously narrowing the search space for structural motifs and broadening our ability to empirically validate results in simulation.

These challenges will likely not be surmounted by individual research teams. Modern neuroscience research requires large cooperative teams with broad-ranging skills and domains of knowledge [40, 41, 1]. In the spirit of this belief, we have open-sourced all of the code, data, and infrastructure developed in the course of this research, including those forays which led to dead ends. We intend for the community to benefit from and apply these resources to their own datasets and research in the future.

## Materials and Methods

We share a greedy motif discovery algorithm, and then discuss our implementation of subgraph search, bias corrections, and evaluation of significance. We also outline our software and compute infrastructure approach, and share resources for the community to replicate our results.

### An approach for iterative subgraph downselection

Our implementation takes the form of a greedy “funnel” to rapidly downselect candidate subgraphs (**Fig. 5**). This pipeline alternates between subgraph downselection phases (where the total search space is reduced) and subgraph significance computations (where each subgraph is assigned a sortable value to determine its relative importance). The process begins with an exhaustive enumeration and search for all undirected subgraphs of a certain size. Our experiments use a subgraph size of |*V*| ≤ 5, and we calibrated our software to run on consumer hardware for small host graphs, of the scale and density of the *C. elegans* synaptic connectome. We then generate a population of randomized host graphs using the *X*-swap technique (see **A bias-corrected *X*-swap operation**), and search within these perturbed host graphs for the same subgraphs as in the original host. This yields a population distribution against which the true subgraph counts may be compared (see **Significance tests**). These subgraphs are then sorted by their significance values — the most significant may be considered undirected motifs for the given connectome. If the user wishes to find directed graphs rather than undirected graphs, a user-specified number of undirected motifs (the top *n* most-significant) then progress to the directed search stage.

**Figure 5.**
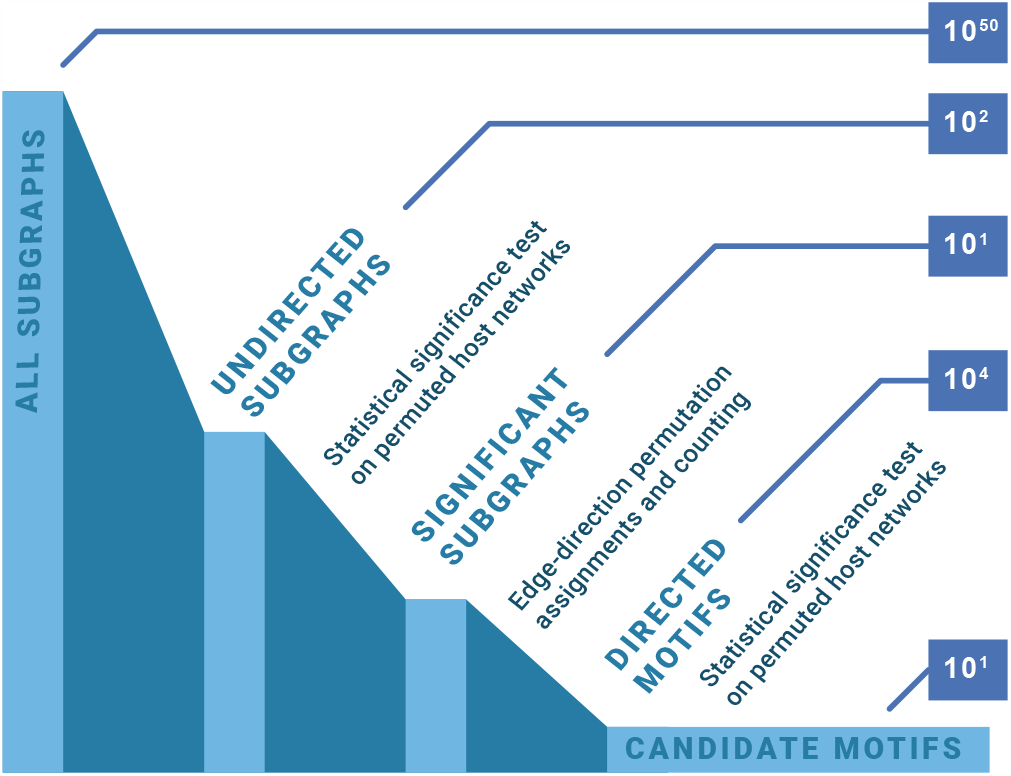
An illustration of the processing pipeline. As an illustration, we provide the rough order of magnitude of the equivalent search space if an exhaustive subgraph search were performed at each stage. Numbers are provided for five-vertex subgraphs. We begin with the set of all possible (directed) subgraphs — a prohibitively large set which cannot be explored exhaustively. An exhaustive search of all undirected subgraphs (only a few hundred) is feasible on consumer hardware. A significance value can then be assigned to each subgraph in this population. A small number of hand-selected subgraphs (*O*(10)) will then be passed onward for edge direction assignment. The total number of possible directed subgraphs that template “onto” these undirected motifs will vary depending on the undirected motifs selected, but will tend to be around *O*(10^4^) *− O*(10^5^). An exhaustive search of this scale is feasible, though may require specialized high-throughput infrastructure for larger host graphs. Finally, significance values are assigned to each directed result subgraph, and a small selection of directed motifs can be returned for further evaluation and study.

### Assigning edge direction to undirected templates

A single undirected graph “template” can yield many connected, directed graphs (**Fig. 6**). In other words, any undirected (symmetric) adjacency matrix *A* can yield a large set of directed matrices *D*, defined by *A*_*D*_ ∈ {*D* such that 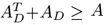 and 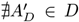 where 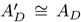} (where (+) and (≥) are element-wise operators). Finding unique, directed permutations of an undirected template graph with unlabeled vertices is not a simple operation [42]. We can place a weak upper bound on the number of possible permutations | *D* | *≤* 4^|*E*|^, since for each (unordered) pair of vertices (*u, v)* in the undirected graph there are four options: no edge, an edge from *u* to *v*, an edge from *v* to *u*, or a bidirectional edge (both (*u, v*) and (*v, u*) exist). This satisfies the first constraint 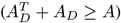 but there is still a large possible number of isomorphic graphs in this set.

**Figure 6.**
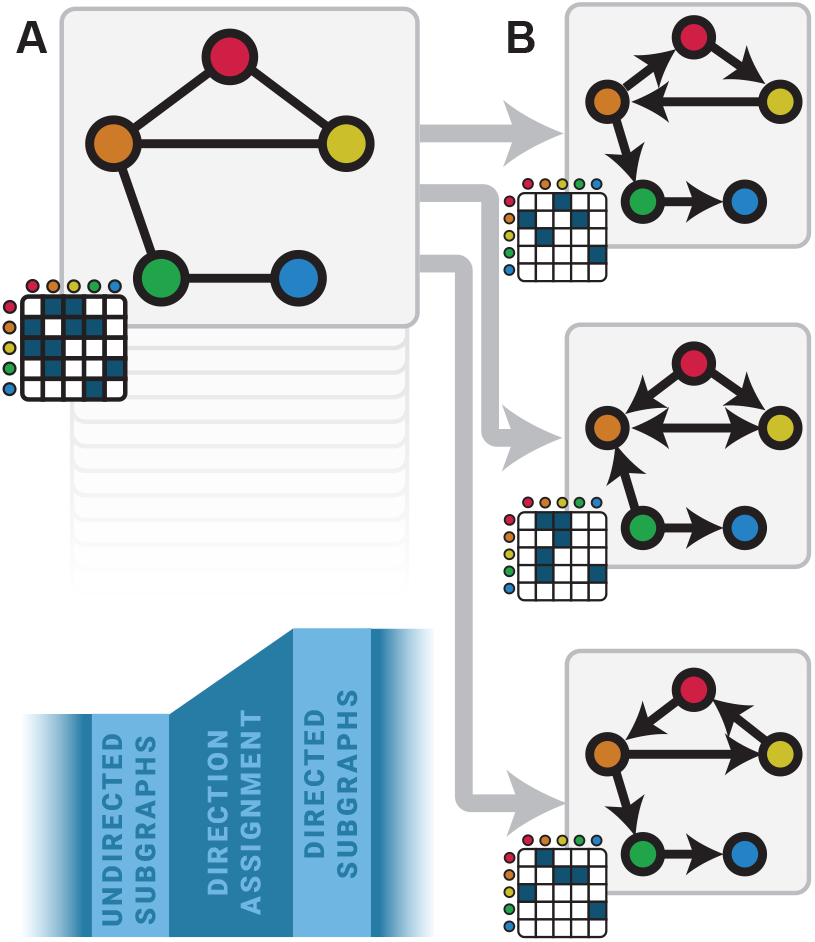
An illustration of the challenges of the edge permutation and direction assignment subprocess of our pipeline (blue inset). **A**. The “template” undirected graph comprises five vertices and five undirected edges. Its adjacency matrix is rendered in the lower left corner. By assigning a direction (*u →v, u ←v*, or both *u↔ v*) to each template edge, we can construct up to 3^5^ possible directed graphs. **B**. Three of those directed graphs are shown here. Though the adjacency matrices are distinct, the top and bottom graphs are isomorphic, and only one should be included in the set of evaluated graphs.

The only way to distinguish unique directed graphs from isomorphic repeats, to the best of our knowledge, is to exhaustively check each. For this reason, it makes sense to keep track of a cache of these graph identities. We compute a base-4 identifier for each graph in the directed set (**Equation 1**).

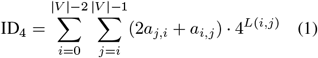

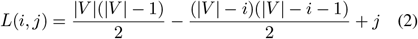

In this formula, |V| represents the vertex count, and *a*_*i*,*j*_ represents the value in the cell (*i, j*) of the adjacency matrix. The cell index (*i, j*) is converted to a linear index of the upper triangle of the square matrix, including the diagonal, using *L*(*i, j*) from **Equation 2** [43]. The double summation 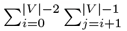 takes into account the upper triangle of the matrix, including the diagonal. (To exclude loops from this identification process, *j* can be redefined as *j* = *i* + 1.) Thus, for each (vertex-labeled) directed graph, we compute a unique ID. Isomorphism mappings between graph IDs can then be cached to avoid recomputing. Our Python library stores a copy of unique graph IDs for all directed graphs with up to five vertices (around ten thousand IDs), as well as the isomorphism mapping to avoid searching for duplicate graphs.

### A bias-corrected *X*-swap operation

The *X*-swap graph randomization technique was first described in [31] as a degree-preserving randomization technique for undirected graphs (**Fig. 1C**). Extending this technique to *directed* graphs is not trivial, as repeated application of the “traditional” *X*-swap operation on a directed graph will result in a nonergodic and biased walk [32] as can be seen in **Fig. 1D** where even an Erdős-Rényi network is found to have “significant” motifs if the plain *X*-swap Monte Carlo operation is performed. Instead, we must also include the option for a *triangle-swap* operation, in which four vertices, randomly selected with replacement, are triangle-swapped if any vertex is chosen twice. The triangle swap involves simply reversing the direction of the edges in the triangle induced by those three vertices, whether or not they induce a directed cycle (**Fig. 1E**). This correction accommodates a graph property known as *mobility*, defined in [32] as

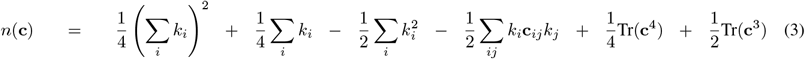

Though computing mobility on even a large host graph can be made relatively efficient through deployment of linear algebra libraries on the GPU, it is even easier to redesign our swap operation so that we do not change *n*(**c**) to begin with, thereby obviating the need to recompute it after each swap. The result is that we now have an unbiased, repeatable *X*-swap operation that runs nearly as fast as the original, from which we can sample many new host graphs in the local neighborhood of our *N* = 1 host. We validate this modified *X*-swap by repeating our earlier experiment on an Erdős-Rényi host network. This time, it yields *no* significant results, as expected.

Prior literature disagrees on how many swap operations need to be run on a graph in order for it to converge on a usefully-distinct state [32]. To calibrate this value for our needs, we defined a *swap-ratio*, or the relationship of swap operations performed to the total edge count of the graph. Counter-intuitively, a swap ratio of 1.0 tends not to be sufficient to fully randomize a graph. For each host graph, we swept the swap ratio hyperparameter across a wide range of values, from 0.0 to 4.0. We found that a swap ratio of 2.0 (2 |*E*| total swaps) tended to result in a well-randomized network for graphs of the scale examined here (**Fig. 7A**). Because the computational cost of the randomization Monte Carlo is linear with the number of swaps (**Fig. 7B**), a swap ratio of 2.0 struck a balance between thoroughness and computational economy.

**Figure 7.**
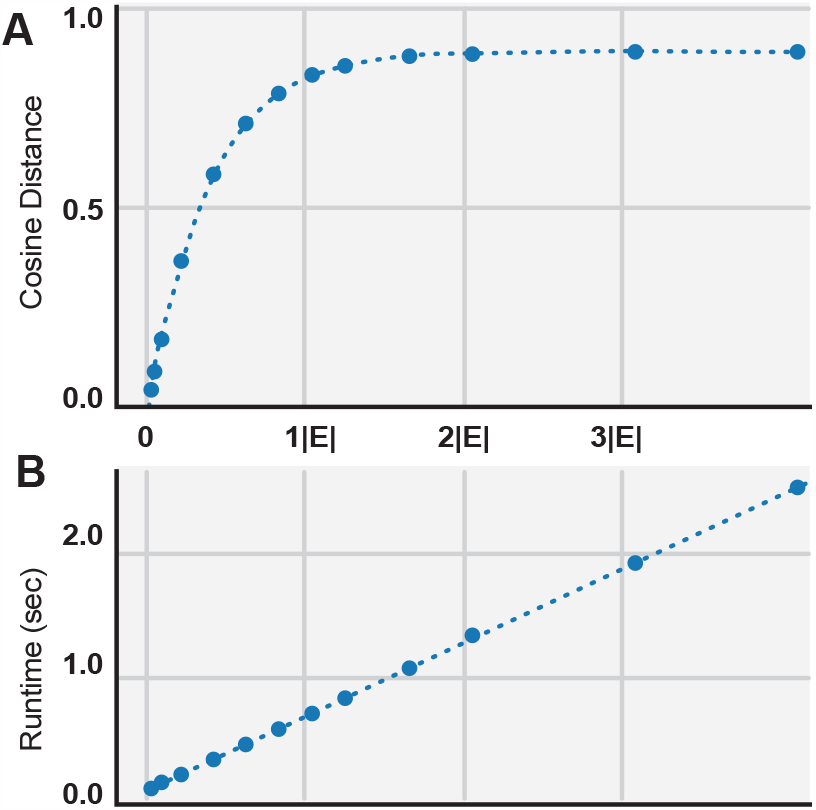
Evaluating the *X*-swap Monte Carlo operation. **A. Parameter-sweep of swap ratio**. Swap ratio is defined as the ratio between the number of *X*-swap operations performed and the total number of edges |*E*| in the host graph. Here, we picture the parameter sweep results for the *C. elegans* connectome, with swap count on the *x* axis and cosine distance on the *y* axis. A swap ratio of 1.0 does not lead to a maximally randomized graph (since edges are selected with replacement prior to each swap and so the same edges may be selected multiple times). We found a swap ratio of 2.0 was more than sufficient to produce a graph with the same degree distribution as the original, but with sufficiently permuted edges. **B. Monte Carlo execution time**. Because the time-cost of the randomization technique is linear with the number of swaps performed, a balance must be struck between swap ratio and execution time.

Crucially, the *X*-swap operation can still affect motif counts as desired: Repeated swaps in both directed and undirected graphs can break or contribute structure (**Fig. 1B-C**) in a way that helps us understand which types of subgraphs arise naturally in a graph with our properties — i.e., which subgraphs exist just as frequently in our null model population — and which subgraphs are “intentional” and occur in our original host network with an unexpected frequency or rarity.

### A test for subgraph significance

After counting subgraphs with an off-the-shelf tool such as GrandIso [10] in both original host as well as a population of randomized graphs, we must now determine a *significance value* for each subgraph. This is most easily done with a traditional one-sample *T*-test, comparing the population statistics to the true motif count (**Fig. 1D**). The resulting statistic gives us a scalar value for each subgraph. This can be used to either sort the subgraphs or threshold to produce a subset. In our undirected pipeline stage, we use a Bonferroni-corrected threshold of 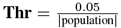 to determine which motifs to promote for further study in the directed stage of the pipeline. Our *C. elegans* and EB studies both used a population size of 10.

### Compute infrastructure

For small host graphs such as the *C. elegans* complete synaptic connectome, we determined that our pipeline can be run in a handful of hours on consumer-grade compute equipment (<1 day on a 2018 Intel MacBook Pro laptop, with 64 GB RAM and eight cores). For larger host graphs and for our parameter sweeping operations, we used an on-premise SLURM compute cluster, for which our job engineering code is available. The much more computationally expensive ellipsoid body graph has a density of 0.389, roughly an order of magnitude higher than *C. elegans*, 0.044 [10, 16]. The ellipsoid body graph took a total of 1,125 CPU-days to run on the compute cluster (around 36,000 searches at *µ* = 45 minutes each). This runtime is dominated by the edge count and local topology of the host graph, rather than its vertex count, and so to our knowledge there is no straightforward way to estimate runtime prior to execution at this time [15]. To further reduce the wallclock execution time, we also provide a cloud-based software implementation that uses serverless compute (i.e., Amazon Web Services Lambda and DynamoDB) [15]. All results from both of these connectome graph searches, as well as our randomized network results, will be made publicly available.

## Acknowledgements

The authors gratefully acknowledge internal financial support from the Johns Hopkins University Applied Physics Laboratory’s Independent Research & Development Program for funding portions of this work. Research reported in this publication was also supported in part by the National Institutes of Health under award number R24MH114785 and R01MH12668. The content is solely the responsibility of the authors and does not necessarily represent the official views of the National Institutes of Health.

## Notes

https://github.com/aplbrain

### Competing Interest Statement

The authors have declared no competing interest.

## References

[1] MICrONS Consortium, J Alexander Bae, Mahaly Baptiste, Caitlyn A Bishop, Agnes L Bodor, Derrick Brittain, JoAnn Buchanan, Daniel J Bumbarger, Manuel A Castro, Brendan Celii, et al. Functional connectomics spanning multiple areas of mouse visual cortex. BioRxiv, pages 2021–07, 2021.

[2] Alexander Shapson-Coe, Michał Januszewski, Daniel R Berger, Art Pope, Yuelong Wu, Tim Blakely, Richard L Schalek, Peter H Li, Shuohong Wang, Jeremy Maitin-Shepard, et al. A connectomic study of a petascale fragment of human cerebral cortex. BioRxiv, pages 2021–05, 2021.

[3] Louis K Scheffer, C Shan Xu, Michal Januszewski, Zhiyuan Lu, Shin-ya Takemura, Kenneth J Hayworth, Gary B Huang, Kazunori Shinomiya, Jeremy Maitlin-Shepard, Stuart Berg, et al. A connectome and analysis of the adult drosophila central brain. Elife, 9:e57443, 2020.

[4] Todd Waskiewicz. Friend of a friend influence in terrorist social networks. In Proceedings on the international conference on artificial intelligence (ICAI), page 1. The Steering Committee of The World Congress in Computer Science, Computer …, 2012.

[5] Vernon B Mountcastle. Modality and topographic properties of single neurons of cat’s somatic sensory cortex. Journal of neurophysiology, 20(4):408–434, 1957.

[6] Daniel Witvliet, Ben Mulcahy, James K Mitchell, Yaron Meirovitch, Daniel R Berger, Yuelong Wu, Yufang Liu, Wan Xian Koh, Rajeev Parvathala, Douglas Holmyard, et al. Connectomes across development reveal principles of brain maturation. Nature, 596(7871):257–261, 2021.

[7] Erik C Johnson, Brian S Robinson, Gautam K Vallabha, Justin Joyce, Jordan K Matelsky, Raphael Norman-Tenazas, Isaac Western, Marisel Villafañe-Delgado, Martha Cervantes, Michael S Robinette, et al. Exploiting large neuroimaging datasets to create connectome-constrained approaches for more robust, efficient, and adaptable artificial intelligence. In Artificial Intelligence and Machine Learning for Multi-Domain Operations Applications V, volume 12538, pages 394–405. SPIE, 2023.

[8] Anthony Zador, Sean Escola, Blake Richards, Bence Ölveczky, Yoshua Bengio, Kwabena Boahen, Matthew Botvinick, Dmitri Chklovskii, Anne Churchland, Claudia Clopath, et al. Catalyzing next-generation artificial intelligence through neuroai. Nature communications, 14(1):1597, 2023.

[9] Dhireesha Kudithipudi, Mario Aguilar-Simon, Jonathan Babb, Maxim Bazhenov, Douglas Blackiston, Josh Bongard, Andrew P Brna, Suraj Chakravarthi Raja, Nick Cheney, Jeff Clune, et al. Biological underpinnings for lifelong learning machines. Nature Machine Intelligence, 4(3):196–210, 2022.

[10] Jordan K. Matelsky, Elizabeth P. Reilly, Erik C. Johnson, Jennifer Stiso, Danielle S. Bassett, Brock A. Wester, and William Gray-Roncal. DotMotif: an open-source tool for connectome subgraph isomorphism search and graph queries. Scientific Reports, 11(1), Jun 2021.

[11] Vincenzo Bonnici, Rosalba Giugno, Alfredo Pulvirenti, Dennis Shasha, and Alfredo Ferro. A subgraph isomorphism algorithm and its application to biochemical data. BMC bioinformatics, 14(7):1–13, 2013.

[12] L. P. Cordella, P. Foggia, C. Sansone, and M. Vento. A (sub)graph isomorphism algorithm for matching large graphs. IEEE Transactions on Pattern Analysis and Machine Intelligence, 26(10):1367–1372, October 2004.

[13] Olaf Sporns and Rolf Kötter. Motifs in Brain Networks. PLOS Biology, 2(11):e369, October 2004.

[14] Shin-ya Takemura, Aljoscha Nern, Dmitri B Chklovskii, Louis K Scheffer, Gerald M Rubin, and Ian A Meinertzhagen. The comprehensive connectome of a neural substrate for ‘ON’ motion detection in Drosophila. Elife, 6:e24394, 2017.

[15] Jordan K Matelsky, Erik C Johnson, Brock Wester, and William R Gray-Roncal. Scalable graph analysis tools for the connectomics community. bioRxiv, 2022.

[16] Jordan K Matelsky, Raphael Norman-Tenazas, Felicia Davenport, Elizabeth P Reilly, and William Gray-Roncal. Circuit motifs and graph properties of connectome development in c. elegans. bioRxiv, 2021.

[17] Alexander B Kunin, Jiahao Guo, Kevin E Bassler, Xaq Pitkow, and Krešimir Josić. Hierarchical modular structure of the drosophila connectome. Journal of Neuroscience, 43(37):6384–6400, 2023.

[18] Albert Lin, Runzhe Yang, Sven Dorkenwald, Arie Matsliah, Amy R Sterling, Philipp Schlegel, Szi-chieh Yu, Claire E McKellar, Marta Costa, Katharina Eichler, et al. Network statistics of the whole-brain connectome of drosophila. bioRxiv, pages 2023–07, 2023.

[19] Brian Matejek, Donglai Wei, Tianyi Chen, Charalampos E Tsourakakis, Michael Mitzenmacher, and Hanspeter Pfister. Edge-colored directed subgraph enumeration on the connectome. Scientific Reports, 12(1):11349, 2022.

[20] Elisabeth Wong, Brittany Baur, Saad Quader, and Chun-Hsi Huang. Biological network motif detection: principles and practice. Briefings in bioinformatics, 13(2):202–215, 2012.

[21] Falk Schreiber and Henning Schwöbbermeyer. Frequency concepts and pattern detection for the analysis of motifs in networks. In Transactions on computational systems biology III, pages 89–104. Springer, 2005.

[22] Falk Schreiber and Henning Schwöbbermeyer. Mavisto: a tool for the exploration of network motifs. Bioinformatics, 21(17):3572–3574, 2005.

[23] Sebastian Wernicke and Florian Rasche. Fanmod: a tool for fast network motif detection. Bioinformatics, 22(9):1152–1153, 2006.

[24] Saeed Omidi, Falk Schreiber, and Ali Masoudi-Nejad. Moda: an efficient algorithm for network motif discovery in biological networks. Genes & genetic systems, 84(5):385–395, 2009.

[25] Ilan Y Smoly, Eugene Lerman, Michal Ziv-Ukelson, and Esti Yeger-Lotem. Motifnet: a web-server for network motif analysis. Bioinformatics, 33(12):1907–1909, 2017.

[26] Michael Winding, Benjamin D Pedigo, Christopher L Barnes, Heather G Patsolic, Youngser Park, Tom Kazimiers, Akira Fushiki, Ingrid V Andrade, Avinash Khandelwal, Javier Valdes-Aleman, et al. The connectome of an insect brain. Science, 379(6636):eadd9330, 2023.

[27] Paul Erdös and Albert Rényi. On Random Graphs I. Publicationes Mathematicae Debrecen, 6:290, 1959.

[28] D.M.S.M. Penrose, M. Penrose, and Oxford University Press. Random Geometric Graphs. Oxford studies in probability. Oxford University Press, 2003.

[29] Réka Albert and Albert-László Barabási. Statistical mechanics of complex networks. Reviews of Modern Physics, 74(1):47–97, January 2002.

[30] Duncan J. Watts and Steven H. Strogatz. Collective dynamics of “small-world” networks. Nature, 393(6684):440–442, June 1998.

[31] Ekaterina S. Roberts and Anthony C. C. Coolen. Unbiased degree-preserving randomisation of directed binary networks, 2011.

[32] Antoon CC Coolen, Andrea De Martino, and Alessia Annibale. Constrained markovian dynamics of random graphs. Journal of Statistical Physics, 136(6):1035–1067, 2009.

[33] John G White, Eileen Southgate, J Nichol Thomson, Sydney Brenner, et al. The structure of the nervous system of the nematode caenorhabditis elegans. Philos Trans R Soc Lond B Biol Sci, 314(1165):1–340, 1986.

[34] Daniel B Turner-Evans, Kristopher T Jensen, Saba Ali, Tyler Paterson, Arlo Sheridan, Robert P Ray, Tanya Wolff, J Scott Lauritzen, Gerald M Rubin, Davi D Bock, et al. The neuroanatomical ultrastructure and function of a biological ring attractor. Neuron, 108(1):145–163, 2020.

[35] Raphael Norman-Tenazas, Brian S Robinson, Justin Joyce, Isaac Western, Erik C Johnson, William Gray-Roncal, and Joan A Hoffmann. Continuous state estimation with synapse-constrained connectivity. In 2022 International Joint Conference on Neural Networks (IJCNN), pages 1–9. IEEE, 2022.

[36] Adriane G Otopalik, Alexander C Sutton, Matthew Banghart, and Eve Marder. When complex neuronal structures may not matter. Elife, 6:e23508, 2017.

[37] Adriane G Otopalik, Marie L Goeritz, Alexander C Sutton, Ted Brookings, Cosmo Guerini, and Eve Marder. Sloppy morphological tuning in identified neurons of the crustacean stomatogastric ganglion. Elife, 6:e22352, 2017.

[38] Caitlyn Parmelee, Samantha Moore, Katherine Morrison, and Carina Curto. Core motifs predict dynamic attractors in combinatorial threshold-linear networks. PloS one, 17(3):e0264456, 2022.

[39] Vladimir Itskov, Carina Curto, Eva Pastalkova, and György Buzsáki. Cell assembly sequences arising from spike threshold adaptation keep track of time in the hippocampus. Journal of Neuroscience, 31(8):2828–2834, 2011.

[40] Larry F Abbott, Davi D Bock, Edward M Callaway, Winfried Denk, Catherine Dulac, Adrienne L Fairhall, Ila Fiete, Kristen M Harris, Moritz Helmstaedter, Viren Jain, et al. The mind of a mouse. Cell, 182(6):1372–1376, 2020.

[41] Joshua T Vogelstein, Eric Perlman, Benjamin Falk, Alex Baden, William Gray Roncal, Vikram Chandrashekhar, Forrest Collman, Sharmishtaa Seshamani, Jesse L Patsolic, Kunal Lillaney, et al. A community-developed open-source computational ecosystem for big neuro data. Nature methods, 15(11):846–847, 2018.

[42] Peter J Cameron. Sequences realized by oligomorphic permutation groups. J. Integer Seq, 3(1), 2000.

[43] Paul Sinclair (https://math.stackexchange.com/users/258282/paul sinclair). Conversion of upper triangle linear index from index on symmetrical array. Mathematics Stack Exchange. URL:https://math.stackexchange.com/q/2134297 (version: 2017-02-08).

